# DNA methylation, combined with RNA sequencing, provide novel insight into molecular classification of chordomas and their microenvironment

**DOI:** 10.1101/2023.05.06.539695

**Authors:** Szymon Baluszek, Paulina Kober, Natalia Rusetska, Michał Wągrodzki, Tomasz Mandat, Jacek Kunicki, Mateusz Bujko

## Abstract

Chordomas are rare tumors of notochord remnants, occurring mainly in the sacrum and skull base. In spite of slow growth, they are highly invasive what makes the treatment challenging. Because of low incidence the molecular background of chordomas is poorly recognized.

Our study aims to determine role of DNA methylation abnormalities in skull base chordomas including its role in deregulation of gene expression. We subjected 32 tumor and 4 normal nucleus pulposus (NP) samples to profiling of DNA methylation with EPIC microarrays and gene expression with RNAseq.

Genome-wide DNA methylation analysis showed two distinct chordoma clusters (subtypes C and I) with different patterns of aberrant DNA methylation. C Chordomas are characterized by general hypomethylation with hypermethylation of CpG islands, while I chordomas are generally hypermethylated. These differences were reflected by distinct distribution of differentially methylated probes (DMPs). Differentially methylated regions were determined in each chordoma subtype indicating aberrant methylation in known tumor-related genes and regions encoding small RNAs in C chordomas. Correlation between methylation and expression was observed in minority of these genes. Upregulation of *TBXT* in chordomas appeared related to lower methylation at tumor-specific DMR in gene promoter.

Gene expression-based clusters of tumor samples did not overlap with DNA methylation subtypes. Nevertheless, the subtypes substantially differ in transcriptomic profile that shows immune activation in I chordomas and enhanced proliferation in C chordomas. Immune enrichment in chordomas I was confirmed with deconvolution methods (cohesively based on methylation and transcriptomic data). Copy number analysis showed higher chromosomal instability in C chordomas. All but one have 9p deletion (*CDKN2A*/*B*) and downregulation of genes encoded in related chromosomal band. No significant difference in patients’ survival was observed between tumor subtypes, however, shorter survival was observed in patients with higher number of copy number alterations.

## Introduction

Chordomas are rare bone tumors originating from remnants of the notochord [1]. They most commonly occur in the sacral spine (about 50%) and in spheno-occipital region (skull base chordomas, about 30%), and the remainder is distributed along the entire length of the spine. These tumors are diagnosed twice as often in men than in women [2]. Chordomas grow slowly, but are highly invasive. Because of the location and growth pattern, complete surgery is commonly unfeasible and relatively high resistance to chemotherapy characterizes these tumors. Consequently, high percentage of local recurrences is observed in chordoma patients [3]. Furthermore, distant metastases of these tumors are also common. The main modality of treatment of skull base chordomas is surgery (mostly transnasal endoscopy) and radiotherapy, which is often used as adjuvant treatment [3].

Due to the low incidence of chordomas the molecular pathogenesis of these tumors remains unclear, although there has been significant progress in that field in recent years. Few important studies on the role of genomic mutations in chordomas were published [4–10]. They revealed genetic changes in known tumor-related suppressors and oncogenes such as *PIK3CA, PTEN* and *CDKN2A* as well as changes in tissue-specific genes such as *TBXT* duplications and protein truncating mutations in *LYST* gene [4,10]. Importantly, the genomic changes were also found in genes encoding proteins involved in epigenetic regulation including recurrent mutations in *PBRM1* and *SETD2* [4,5,10]. Contrary to the role of genetic abnormalities in pathogenesis of chordomas, much less attention was paid to the role of epigenetic changes. Although methods for analyzing DNA methylation profile in humans have been available for years, only very few papers on the genome-wide DNA methylation have been published to date [11–14]. These studies showed changes in chordoma DNA methylation profile in comparison to their normal counterparts [12] and very recently a clinical relevance of genome-wide DNA methylation pattern was shown [13,14].

Some light has also been shed on chordoma transcriptome [15–17]. Some researchers were struggling to identify any subgroups within their populations [15], while others were able to identify two expression clusters [18]. Comparisons with nucleus pulposus – a physiological remnant of the notochord – have revealed differences in expression of brachyury (*T*) [15,16], *SAMD-5* [16], and other genes associated with development [15]. A recent multi-omic study of chordoma cell lines identified *CA-2* and *THNSL2* as potentially druggable genes of interest in chordoma [17].

The aim of our study was to investigate genome-wide DNA methylation changes in skull base chordomas and the relationship between the methylation profile and gene expression. To our knowledge, this is the first publication that integrates DNA methylation and gene expression profiles from chordoma patients.

## Materials and Methods

### Patients and tissue samples

Thirty-two patients with skull base chordoma were included in this study. They were treated with transnasal and/or transoral endoscopic surgery in Department of Neurosurgery, Maria Sklodowska-Curie National Research Institute of Oncology, Warsaw, in years 2014 – 2020.

Each tumor sample was split and one part of the tissue was used for routine diagnostic procedures while the second one was snap frozen in liquid nitrogen and stored for molecular analysis. Histopathological diagnosis of chordoma according to WHO criteria [19] was confirmed for all the patients. All the tumors were diagnosed as classical chordoma. Overall patients characteristic is shown in Table 1.

Four samples of nucleus pulposus (NP) were obtained from intervertebral disks, collected during discectomy of 4 patients, suffering from degenerative lumbar spine disorder. Nucleus pulposus samples were enzymatically digested for 4 hours at 37°C with 0.2% collagenase type II (Sigma–Aldrich) in a serum-free DMEM, based on previously validated protocol [20]. The digested tissue/cell suspension was filtered through sterile nylon fabric to remove remaining tissue debris. The cells were subsequently centrifuged at 300 x g for 5 min ad subjected to DNA/RNA isolation.

The study was approved by the local Ethics Committee of Maria Sklodowska-Curie National Research Institute of Oncology in Warsaw, Poland. Each patient provided informed consent for the use of the tissue samples for scientific purposes.

### Nucleic acid isolation

Genomic DNA and total RNA from tissue samples were isolated using AllPrep DNA/RNA/miRNA Universal Kit (Qiagen). The procedure included tissue homogenizing with rotor stator homogenizer Omni Tissue Master (Omni International). The concentration of samples was measured both spectrophotometrically using NanoDrop 2000 (Thermo Scientific) and with fluorescence-based method using QuantiFluor Dye kit (Promega) and Quantus (Promega) instrument. Isolated total RNA was stored at -80 °C, whereas genomic DNA was stored at -20 °C.

### Genome-wide DNA methylation profiling

DNA from 32 skull base chordomas and 4 NP samples were bisulfite converted with EZ-96 DNA Methylation kit (Zymo Research, Orange, CA, USA) and used for genome-wide DNA methylation profiling with Methylation EPIC (Illumina) BeadChip microarrays. Recommended protocol for Infinium MethylationEPIC Kit was used (Infinium HD Methylation Assay Reference Guide, Illumina). Laboratory procedures were performed by Eurofins Genomics service.

### Gene expression analysis based on whole transcriptome sequencing

Whole-transcriptome expression profile based on RNA sequencing (RNA-seq) was determined for 32 skull base chordoma and 4 NP samples. One μg of total RNA from each tissue sample was used for library preparation with NEBNext Ultra II Directional RNA Library Prep Kit for Illumina. NEBNext rRNA Depletion Kit was applied for ribosomal depletion. The quality of libraries was assessed using the Agilent Bioanalyzer 2100 system (Agilent Technologies, CA, USA). Libraries were then sequenced on an Illumina NovaSeq 6000 platform, and 150-bp paired-end reads were generated. A minimum of 30 M read pairs per sample were generated. Sequencing was performed by Eurofins Genomics service.

### Data analysis

Data were analyzed, utilizing R statistical programming language (version 4.2.2). Code is freely available in a GitHub repository.

DNA methylation was analyzed using minfi [21] - all samples passed quality control. Methylation probes were filtered (SNP, probes that have failed in at least 25% of samples). Top one percent probes with β-value standard deviation above 0.1 were utilized for gaussian mixture modelling-based clustering [PMID:27818791], for cross-validation hierarchical clustering (euclidean distance and ward.D method, base R package) was utilized. Subsequently, probes differentially methylated between clusters, nucleus pulposus, and chordoma were identified using minfi [21]; p-values lower than 9e-8 were deemed significant [22]. The M-values distributions were tested in a linear model and related β-values were visualized with violin plots. Subsequently, differentially methylated regions were identified with comb-p [23].

Reads abundance on gene and transcripts levels were quantified using kallisto [24] on GRCh37 patch 13 cDNA sequences, downloaded from Ensembl genome database (http://grch37.ensembl.org/Homo_sapiens/Info/Index). Differential gene expression and gene expression normalization was performed, utilizing DESeq2 [25]. Clustering of genes most variable, selected by squared coefficient of variation [26], was performed with gaussian mixture modelling-based clustering [27], for cross-validation hierarchical clustering (euclidean distance and ward.D method) was used. Interaction between methylation and gene expression data was analyzed by means of Kendall correlation of averaged M-values within 1500 bps of TSS and scaled gene expression. Methylation-controlled genes were thusly identified. Similar approach (with mean M-values as proxy for DMR methylation level) was utilized for DMRs.

Subsequently, fold-change differences were analyzed utilizing Fast Gene Set Enrichment Analysis [28] with Gene Ontology and Reactome terms, downloaded from MSigDB [29,30]. Furthermore, weighted correlation network analysis (WGCNA) [31] was utilized to obtain modules of genes, whose association with each methylation cluster was tested with U-Mann-Whitney test. Subsequently, selected modules were intersected wit protein-protein interaction database STRING [32]. Importance of genes in the network was inferred using WGCNA connectivity measure and authority centrality [33], provided by tidygraph package [https://tidygraph.data-imaginist.com].

The tumor microenvironment was deconvoluted from DNA methylation and RNA sequencing – utilizing MethylResolver [34] and immunedeconv [https://github.com/omnideconv/immunedeconv], respectively. In immunedeconv two methods – MCPcounter [35] and ESTIMATE [36] were implemented. The latter was also used in order to compare chordoma clusters with the common human cancer types. Subsequently, non-parametric statistical tests were utilized, where appropriate.

DNA copy number changes were inferred in two-fold manner – conumee was run on the methylation data (http://bioconductor.org/packages/conumee/) and gsealm [37] with data for chromosome bands from MsigDB [29,30]. Non-parametric statistical tests were utilized, where appropriate and Cox-hazard risk model was utilized to test for the survival outcomes. All visualizations were performed with ggplot2 R library[https://ggplot2.tidyverse.org].

## Results

### Chordoma genome-wide DNA methylation profile

Following quality control a set of 834,918 array probes (excluding SNP regions and probes with high probe detection p-value) were analyzed in all 32 samples of skull base chordoma and 4 NP samples. Subsequently, top 3648 most variably methylated probes (top one percent of variable probes from a more general set of probes with standard deviation of β-values above 0.1) were clustered with two independent clustering methods (hierarchical clustering and gaussian modelling-based clustering). Results obtained with both methods showed the same three separate clusters - two clusters composed of chordoma samples (chordoma I - 23 samples and chordoma C with 10 samples, clusters naming refer to the previously proposed terms [13]) with the 4 nucleus pulposus samples in the remaining cluster (Figure 1a). Interestingly, results of hierarchical clustering indicate that chordoma I samples were more similar to NP samples than to chordoma C.

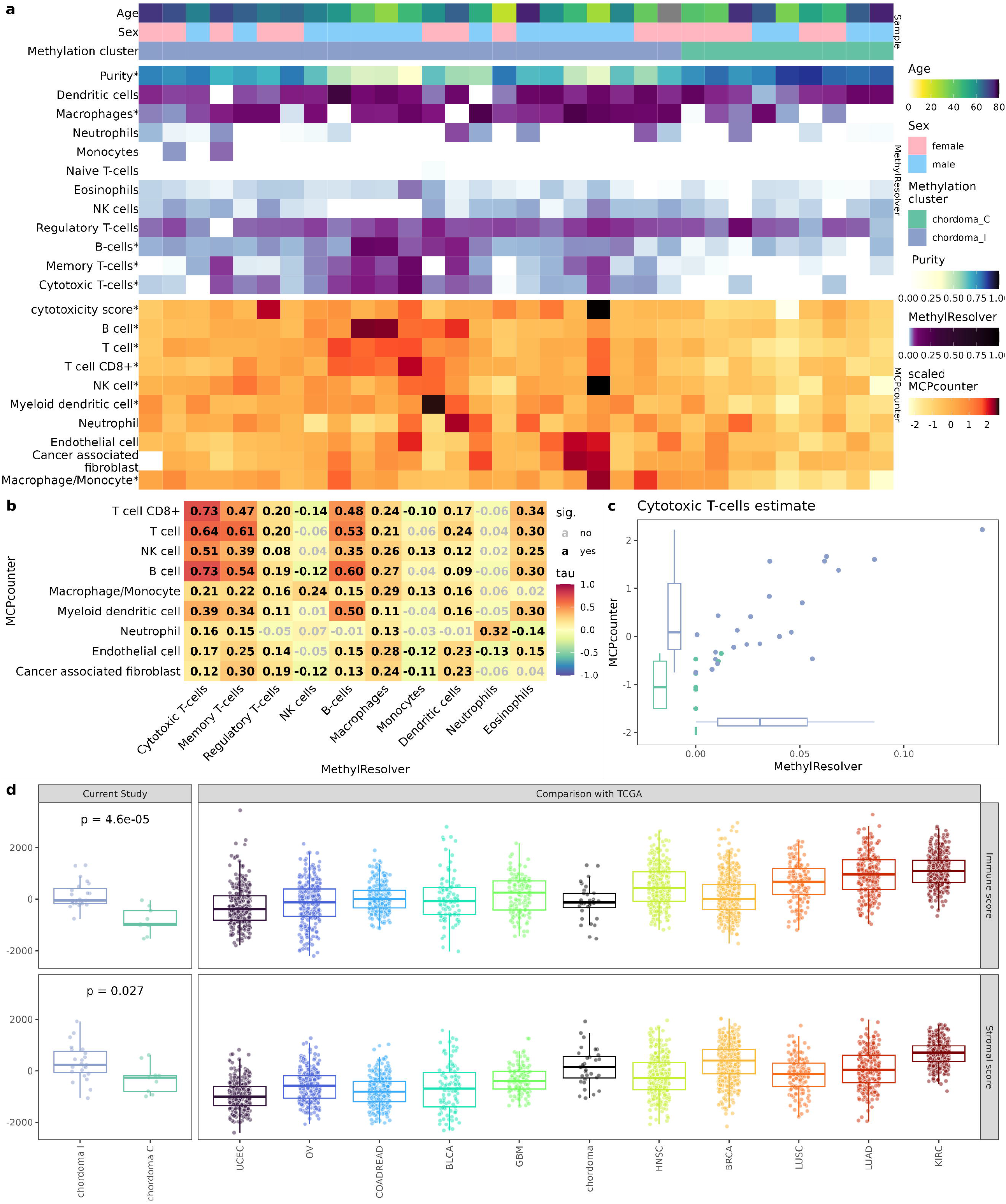

On closer inspection, genome-wide DNA methylation was notably lower in chordoma C than in NP. However, this was most prominent in probes located at Open Sea regions as illustrated by methylation pattern of top variably methylated probes (Figure 1a). This initial observation was further investigated in more general set of variable probes (364,784 probes with standard deviation of β-values above 0.1; Figure 1b,c). This confirms that general hypomethylation in cluster C chordomas was most pronounced in Open Sea. It was less apparent in CpGs closer to CpG islands (CGIs) (including CGI shelves and shores) while the probes in CGIs were hypermethylated in comparison to NP. In turn, chordoma I cluster was globally only slightly hypermethylated; however, this effect became more pronounced in Islands and Shores (Figure 1b). When relation to gene was considered, a more nuanced image emerged – chordoma I samples displayed clear hypermethylation in promoters, chordoma C samples varying levels of hypomethylation. In both gene bodies and intergenic regions, chordomas from C cluster were clearly hypomethylated (Figure 1b). In order to quantify those observations a linear model on M-values was built. Its results, depicted in Figure 1c, warrant explanation: e.g. methylation levels were compared with nucleus pulposus using Open Sea samples/probes as reference. Based only on that, hypomethylation in chordoma C and islands would be expected. However, as described earlier, this was not the case, and therefore effect estimation for combination of chordoma C and Island was strongly positive. As chordoma I samples were generally more methylated, this combination estimate had lower value. Similar but slightly weaker effect was seen in probes relation to genes – probes located on 5’ end (5’UTR, TSS1500, TSS200 and 1^st^ exon) were globally hypomethylated, as were chordoma C samples. However, combination effect was positive, resulting in methylation levels comparable to NP in these 5’ end probes. Besides providing formal evidence to the previously made claims, this model demonstrates that similar hypermethylation of promoters, relative to the intergenic region occurred in chordoma C samples, however, the effect size was smaller and consequently globally hypomethylated pattern dominates at promoters in this cluster.

Subsequently, differentially methylated probes (DMPs) between two chordoma clusters, as well as between chordomas and NP were identified (listed in Supplementary Table 1, summarized in Table 2). Accordingly with results showing higher global DNA methylation in chordoma I than chordoma C subtype, vast majority (9 to 4892) of DMPs identified in comparison of two subtypes of tumors were the probes hypermethylated in I chordomas.

Closer inspection of DMPs character is possible in Figure 1d – even though chordoma C hypermethylation was most pronounced in the promoters and islands, majority of DMPs was found in the Open Sea and gene bodies.

This probably can be explained by the general distribution of probes in the EPIC array and discrepancy in cluster size (as cluster I is bigger, more probes, differing in chordoma versus NP comparison, pass the significance threshold). Furthermore, general pattern of global hypermethylation of chordoma I is reflected by the number of hypermethylated probes in chordoma C in comparison with nucleus pulposus. Proportions of DMPs annotated to particular genomic location categories are shown in Figure 1d.

The DMPs were aggregated into differentially methylated regions (Figure 1e and Supplementary Interactive File 1, which allows reader to interactively explore all DMRs). There were far more hypermethylated DMRs in chordoma I than chordoma C samples both in direct comparison of two subtypes of the tumors and in comparison of each subtype with NP. DMRs with most significant difference between groups of samples and DMRs located in the neighborhood of known cancer-related genes were marked in Figure 1e. Comparison of two chordoma subtypes showed that most significant DMRs are located on chromosome 14 and 19 regions containing microRNA clusters as well as on chromosome 17 region coding for *KRT15, KRT19* and *JUN*. These regions are hypermethylated in chordoma I, when compared to chordoma C.

DMR identified in comparison of chordoma and NP were located at genes with known role in tumor biology, including *TERT, BLM, CDH11, CDH4, DLC1, OPCML, HIF1A, YWHAQ, MGMT, TP63, MTOR, MUPCDH, RIPK4, EGFR* or *TBC1D16*. Most of these DMRs are significantly more hypermethylated in chordoma I (Figure 1e, Supplementary File 1). Chordoma-specific DMRs were also found in location of homeobox domain genes (e.g. DMRs in *HOXA4, HOXA5, HOXD3, HOXD4 MNX1*, and *NFIX*) especially on chromosome 2 at *HOXD* cluster. Moreover, DMR was also identified in *TBXT* (T) that encodes brachyury – a notochord-specific transcription factor that plays developmental role. DMR *in TBXT* was hypomethylated in both chordoma clusters as compared to NP.

### Gene expression in chordoma DNA methylation clusters

Gene expression profile was determined in each chordoma and NP sample based on RNA-seq data. An average 41,315,537 reads per sample were obtained with average 79.65% reads pseudoaligned to UCSC hg19 cDNA transcriptome. The sequencing reads were quantified on 39,293 human transcripts. Of those, 23,013 passed the filtering criteria (at least 5 reads in at least 10 samples) for normalization and differential expression analysis. Number of differentially expressed genes for each condition is available in Table 2.

The results of clustering of the samples based on the expression of most variably expressed genes (4275 genes selected, based on squared coefficient of variation model [26]) did not readily overlap with the results of DNA methylation-based clustering. As a result, one major chordoma expression cluster, including 30 samples, was observed with remaining 3 samples and NP in two remaining separate clusters (Figure 2a). To further investigate the potential relationship between DNA methylation and gene expression entanglement analysis was applied. The result of entanglement 0.22 indicates that, despite general differences between methylation and expression-based clustering, clustering structures are not entirely dissimilar, pointing to more discrete correlations at few-sample level (Supplementary Figure 1). Furthermore, fraction of genes, remaining under methylation control was determined (Figure 2b). Gene was classified as methylation-controlled, when correlation of mean promoter methylation and gene expression was significant and negative, as it is commonly done [38]. In general, 2.9% of genes fulfilled these criteria. However, this fraction was significantly higher in genes differentially expressed in chordoma (Figure 2b). This indicates that different methylation pattern of chordomas influences gene expression on a global scale.

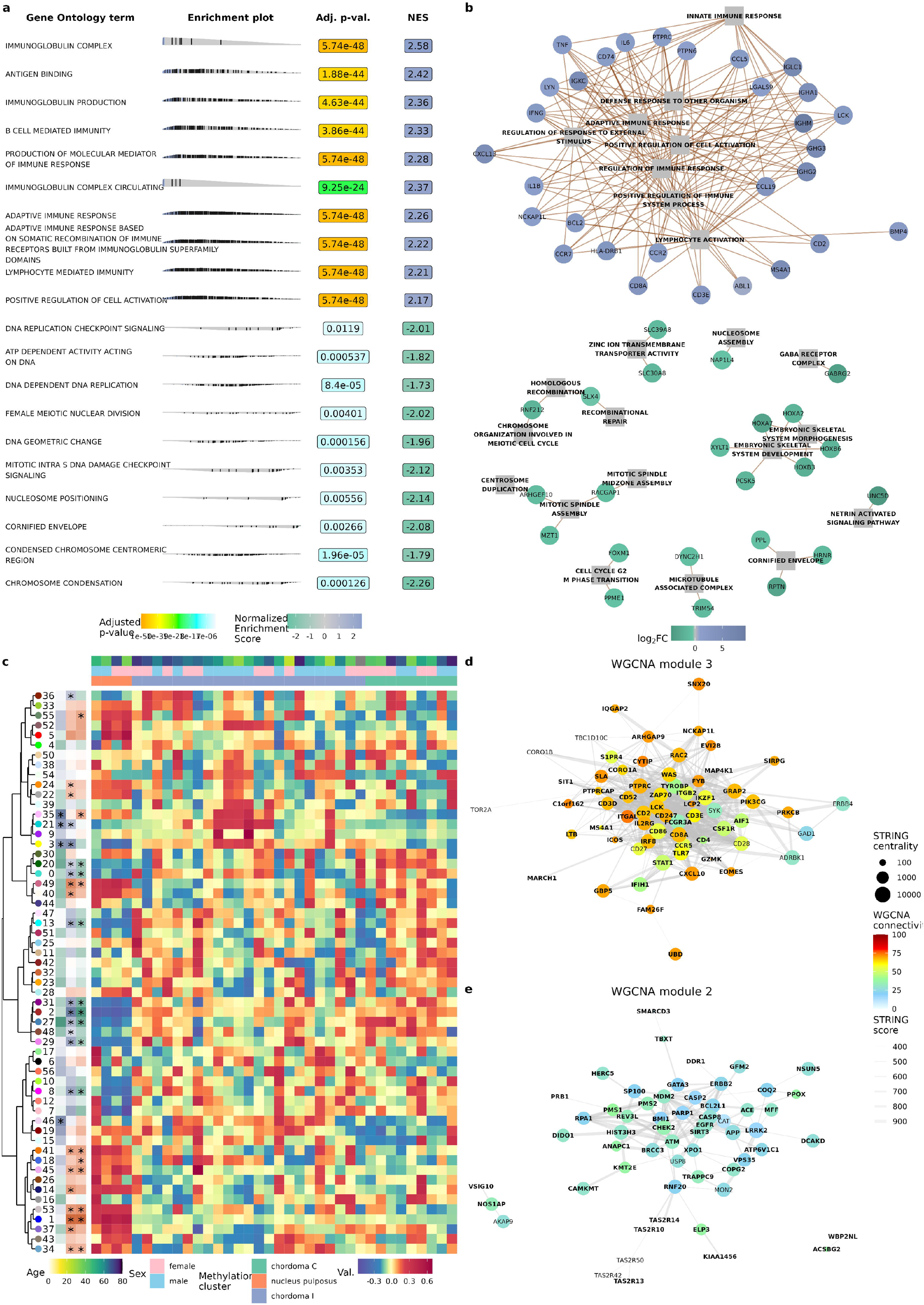

Volcano plots of differentially expressed genes are shown in Figures 2c (chordoma vs nucleus pulposus) and 2d (chordoma I vs chordoma C). Genes marked on the volcano plot were either implicated earlier in chordoma biology (e.g. keratins – *KRT8, KRT18, KRT19, TBXT, LMX1A*, and *EGFR*) or identified by outstanding p-value (e.g. *IL11, CD24*), high absolute fold-change (e.g. *GABRA1*) or DMR result (e.g. *MNX1*). Genes demonstrated on this plot tend to belong to three general categories: immune response-related (e.g. *IL11, ICAM4, CXCL5*), involved in development (e.g. *TBXT, MNX1, HOXA9, SOX9*), and marking epithelial or connective tissue (keratins, *SDC4, PRG4*). In the case of comparing clusters with each other, overlap with existing chordoma cluster markers was less striking – only *KIT* and *CDKN2A* were overexpressed in chordoma I. Other genes, related to this cluster, were skewed towards immune infiltration (HLA proteins, leukocyte markers, cytokines) and high *KIT* expression can be also considered to be related to immune cell signaling. *CDKN2A* loss was previously described in chordoma [39] and evidence of this phenomenon was also seen in further analysis in chordoma C cluster (see section DNA copy number changes in two epigenetic subtypes of chordomas and Figure 5). Furthermore, higher expression levels of homeobox-containing genes were observed in chordoma C.

The relation between DMRs and DEGs was examined. 332 of 1128 genes overexpressed in chordomas I and 42 of 178 genes overexpressed in chordoma C were the ones located in a particular DMR or its close neighborhood, as presented on Figure 2e. Using correlation analysis we determined the genes and DMRs with correlated expression/methylation levels. In summary, 209 such DNA methylation-corelated genes that are differentially expressed and differentially methylated in comparison of two chordoma subtypes were identified. The expression of these 209 genes was correlated with methylation levels of 452 DMRs, what means that expression of more than half of DNA methylation-controlled DEGs were correlated with methylation of more than one DMR. Majority (83%) of these DMRs were hypermethylated in chordomas I and revealed positive methylation/expression correlation. Most of them were located in the gene body thus the result on positive methylation/expression correlation correspond to generally observed role of methylation in gene body regions [40]. This is also in line with observations from DMPs analysis and comparing global DNA methylation levels both indicating that chordomas I had a more hypermethylated epigenetic landscape in general. Top 10 genes with a highest correlation coefficient are listed on the Figure 2e Detail results of correlation analysis are resented in Supplementary Table 7.

Similar analysis was performed for matched DEGs and DMRs that were found in comparison of chordoma vs NP. It showed that expression of 1033 DEGs is correlated corelated with methylation level of total 1543 DMRs. Mainly negative methylation/expression correlation was found as it was observed in case of 75,5 of expression-related DMRs.

For genes with at least moderate correlation between DNA methylation and expression level (correlation coefficient >0.3) we ran overrepresentation analysis with Gene Ontology and Reactome databases to investigate if there are specific pathway enrich for the methylation-controlled DEGs. When analysing methylation-correlated genes that are differentially expressed in chordoma I and C subtypes we found the enrichment of particular terms related basically to immune inflammation and signalling by Rho GTPases. Overrepresentation analysis of DEGs with a corresponding DMR found in comparison of chordoma vs NP showed the enrichment of terms related mainly to extracellular structure organization, cellular junction, epithelial/mesenchymal transition. The significantly enriched terms are listed in Supplementary table 7.

Special attention was paid on to two genes – *TBXT* (brachyury, with official symbol T) and *PTPRCAP*. Brachyury is already recognized as a chordoma marker. Moreover, as presented on Figure 2f, its expression is correlated with methylation in the promoter. CGIs best differentiating chordoma from nucleus pulposus flank the gene promoter and two DMRs were identified up- and downstream to the *TBXT* promoter (Figure 2h, which serves as a genomic map of probes and DMRs). On the other hand, *PTPRCAP* is only one of many immune-related genes, characteristic of chordoma I. It was selected, due to its high promoter methylation-expression correlation (Figure 2g). In fact, all probes, overlapping with the gene were identified as DMRs (Figure 2h).

### Gene set enrichment and weighted correlation network analyses

To characterize functional differences between the two chordoma clusters and nucleus pulposus, gene set enrichment analysis (GSEA) with terms from Gene Ontology (GO, Figure 3a, Supplementary Table 4) and Reactome (Supplementary Table 4) databases was performed. The results displayed striking difference between clusters I and C - genes overexpressed in chordoma I were enriched in terms associated with immune response, with terms associated with adaptive immune response on the very top of the list. Genes sets, characterizing cluster C, were more heterogenous, but pointed to terms related to cell cycle and proliferation with keratinization process, and developmental pathways (Figure 3a). Belongingness of DEGs to the top identified terms is shown on Figure 3b.. Probably stemming from the characteristic of chordoma I cluster (which outnumbered chordoma C cluster 23 to 9), the main difference between chordoma and nucleus pulposus were associated with immune system terms. Nucleus pulposus was enriched in terms associated with cartilage and connective tissue differentiation, possibly pointing to deficiency of those processes in chordoma.

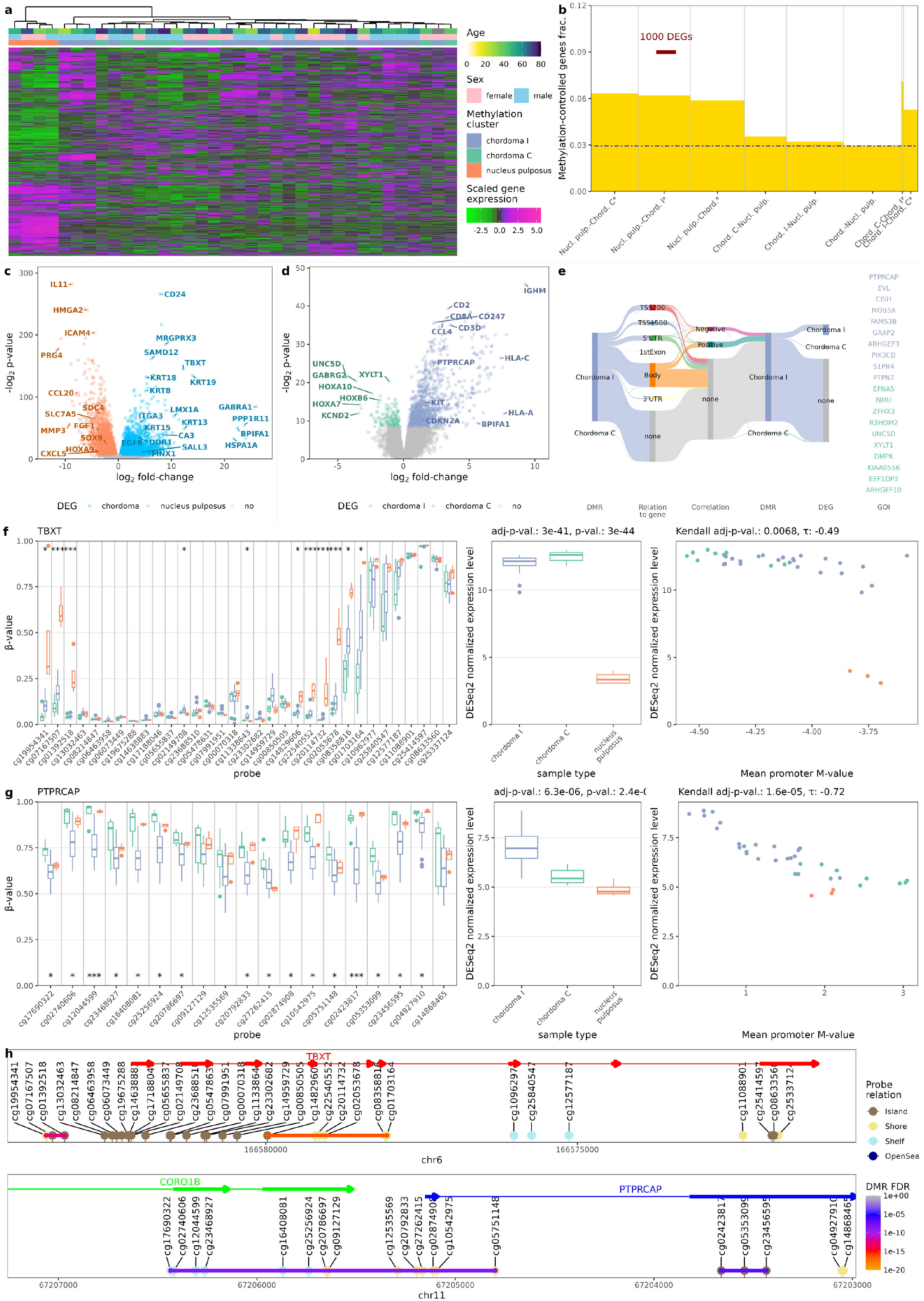

Subsequently, weighted correlation network analysis (WGCNA) was undertaken in order to disentangle gene-gene correlations and operate on fewer, easier to understand modules. Fifty-six such modules were identified (Figure 3c, Supplementary Figure 2). Modules were clustered and differences between chordoma methylation clusters and between each of them and nucleus pulposus were tested (see Table 1 for summary). A cluster of modules 2, 27, 29, 31, and 48 had consistently higher eigenvalues in chordomas while cluster of modules 3, 21, and 35 was associated with chordoma I. Modules 2 and 3 had lowest p-value and log2 fold-change in respective comparisons. In order to validate this finding, clusters were overlapped with protein-protein interactions in STRING database. Notably, *TBXT* was found in module 2 and *PTPRCAP* in module 3. Both were significantly enriched in protein-protein interactions (p < 1e-16 for each). Genes, treated as nodes in network were assigned centrality measures (centrality measures indicate how important is the gene for the network; see methods for details) and WGCNA connectivity scores, which answer a question of how much intra-versus extramodular correlations each gene has. Due to a large number of genes in modules (2690 in module 2 and 2109 in module 3) only genes with high centrality or connectivity scores are shown in Figures 2d and 2e. Genes central in module 3 are immune response-related (e.g. *CD247, CD3E, CD8A, IL2RG*, and *ITGAL*). Contrastingly, genes central for module 2 (and thus probably important for functioning of chordomas in both clusters) are either responsible for cell division and DNA repair (e.g. *ATM, CHEK, BMI1, MDM2*) or parts of inhibitors of apoptosis cascade (e.g. *CASP2, CASP8, SIRT2*).

### Immune infiltration in chordomas

Since GSEA, following differential expression analysis of two chordoma subtypes, clearly indicated difference in the immune response, we made an attempt to estimate the content of immune cells in chordoma samples. Deconvolution methods based on genome-wide DNA methylation data and gene expression profiles were used for this purpose independently. The results of both approaches consistently indicated notably higher content of immune infiltrating cells in chordomas I than in chordomas C. In MethylResolver estimate on DNA methylation chordomas I had higher total immune infiltration (median 0.65 vs median 0.82, p=3.4e-5, Figure 4a). That was reflective of a higher estimated amount of cytotoxic T-cells (p = 1.4e-4, p = 3.0e-4), B-cells (p = 6.4e-4, 4.8e-6) and macrophages (p=0.001, p = 0.006) in both approaches (Figure 4a). A relatively high inter-method scores correlation was observed, i.e. inter-method correlation of abovementioned populations signatures were high (Kendall T 0.73, 0.60, and 0.29 respectively; see Figure 4b). Both methods results for cytotoxic T-cells is additionally shown on Figure 4c.

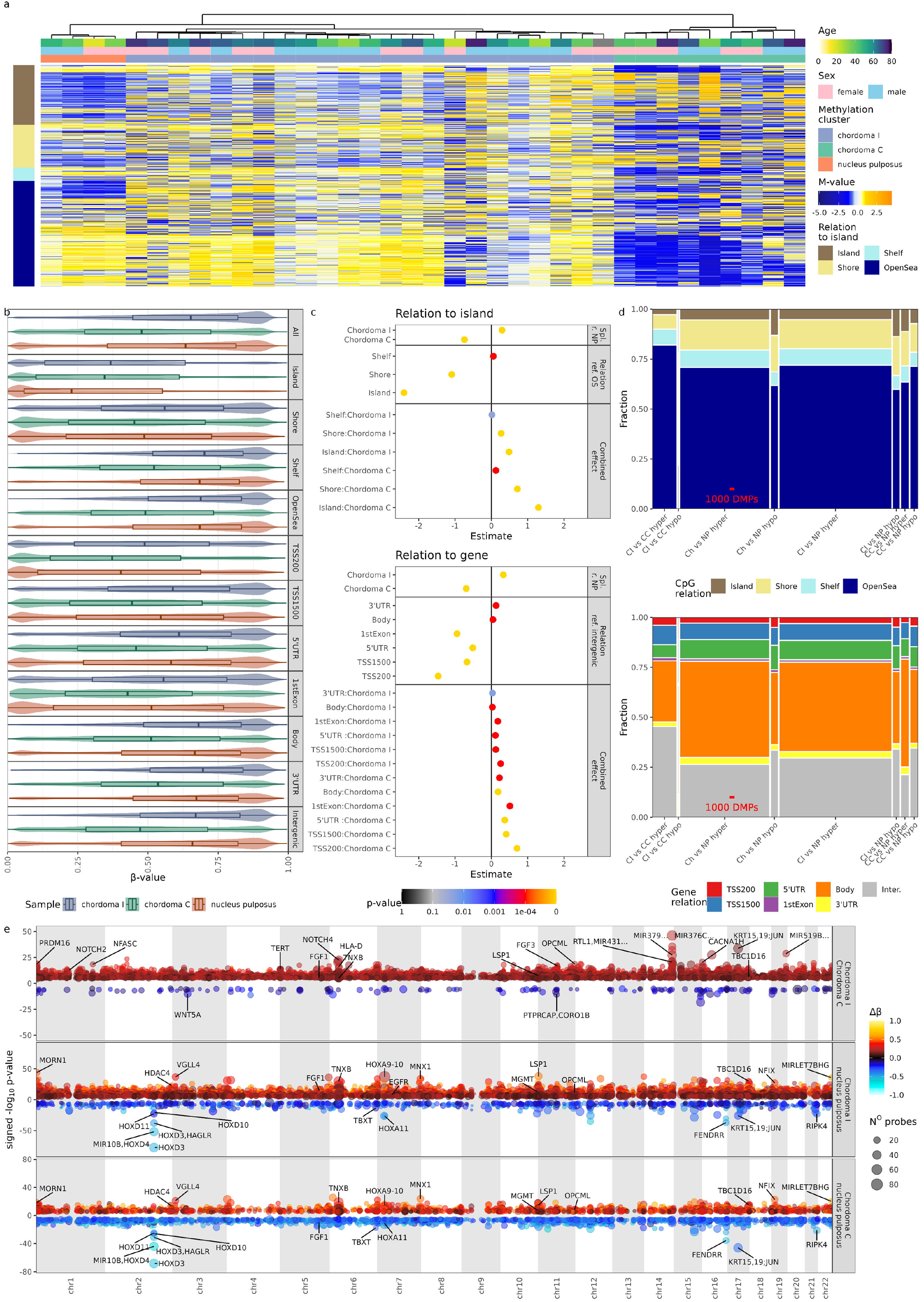

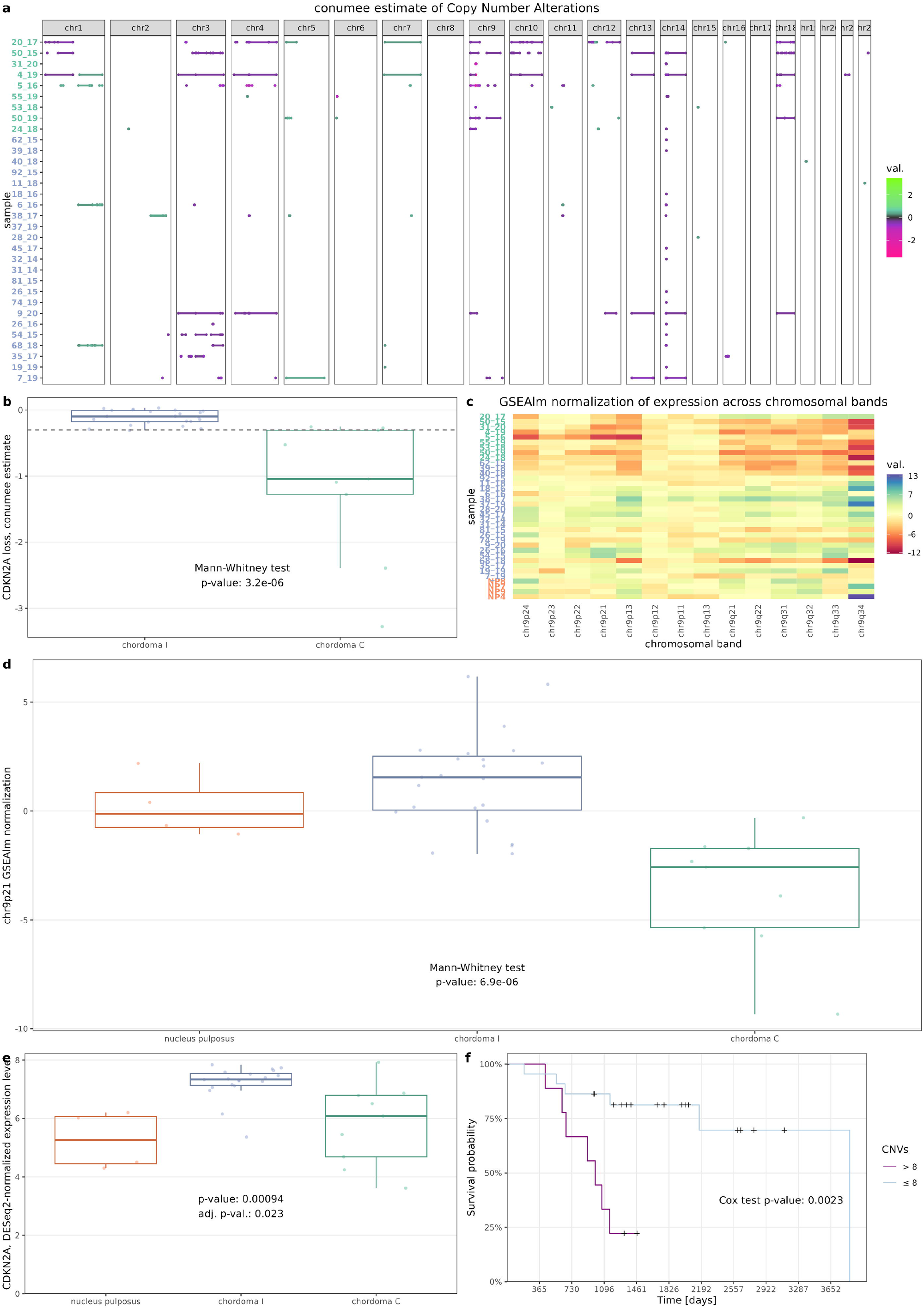

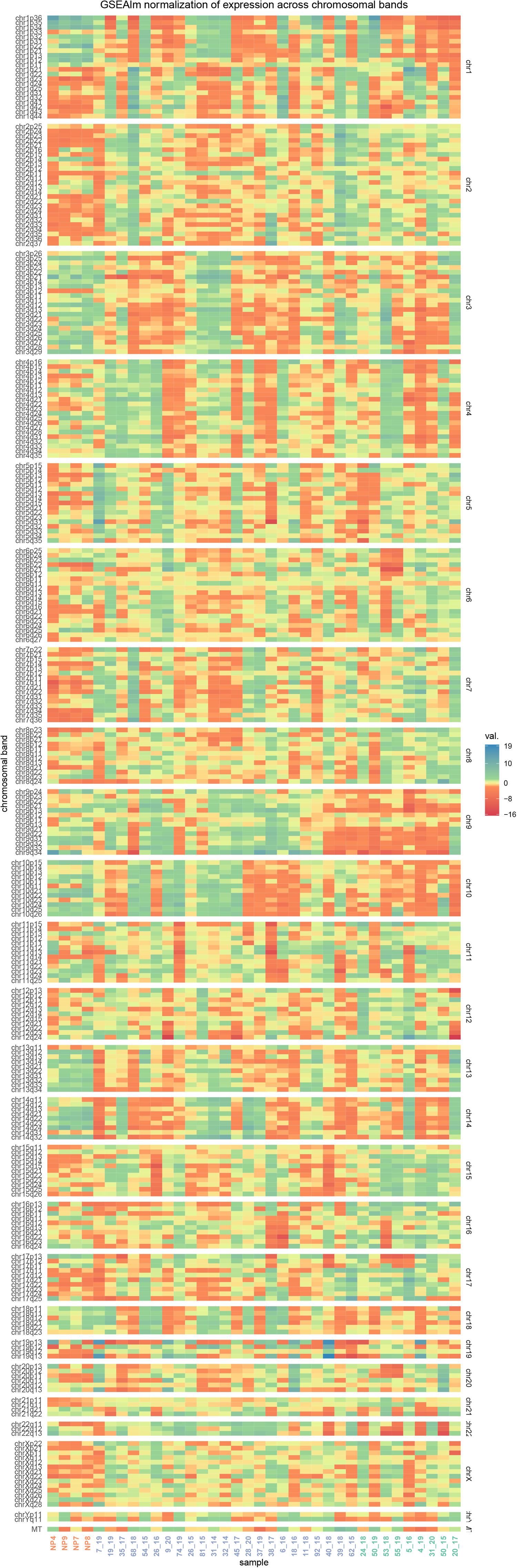

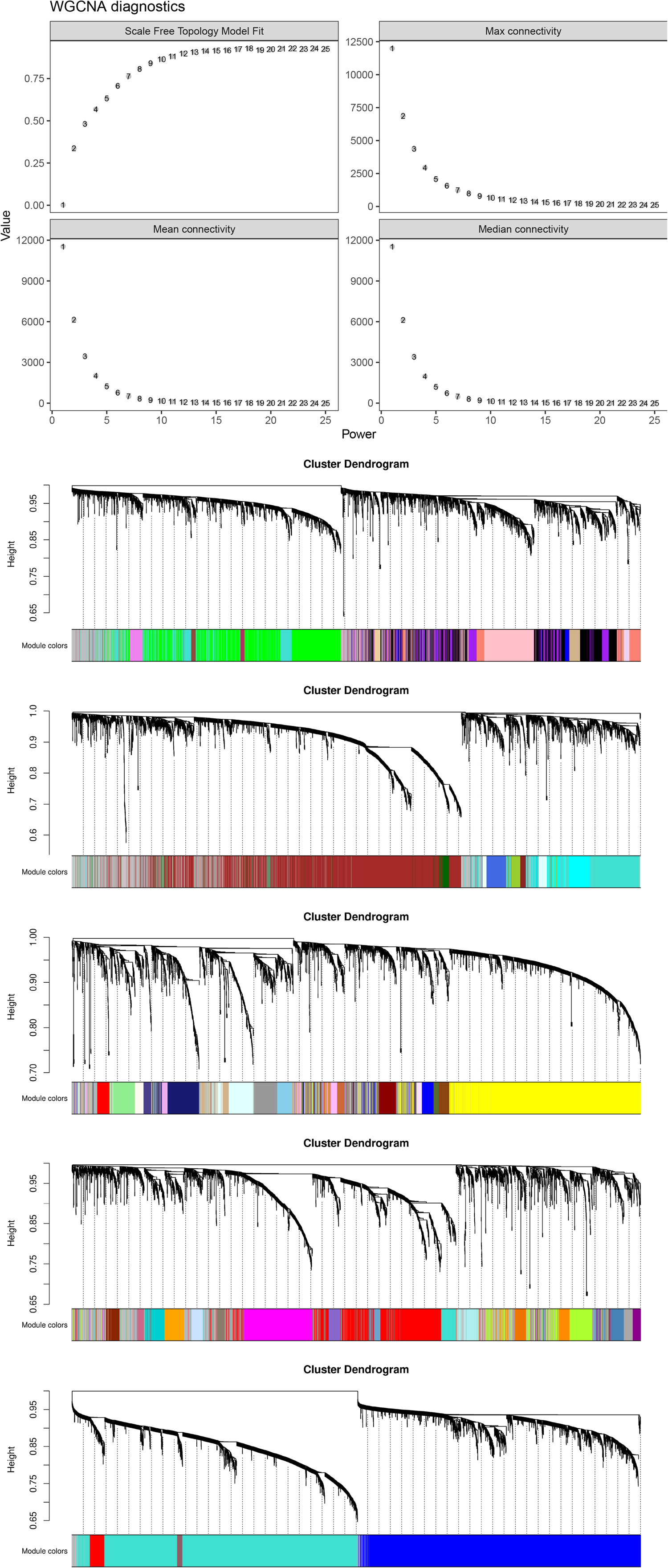

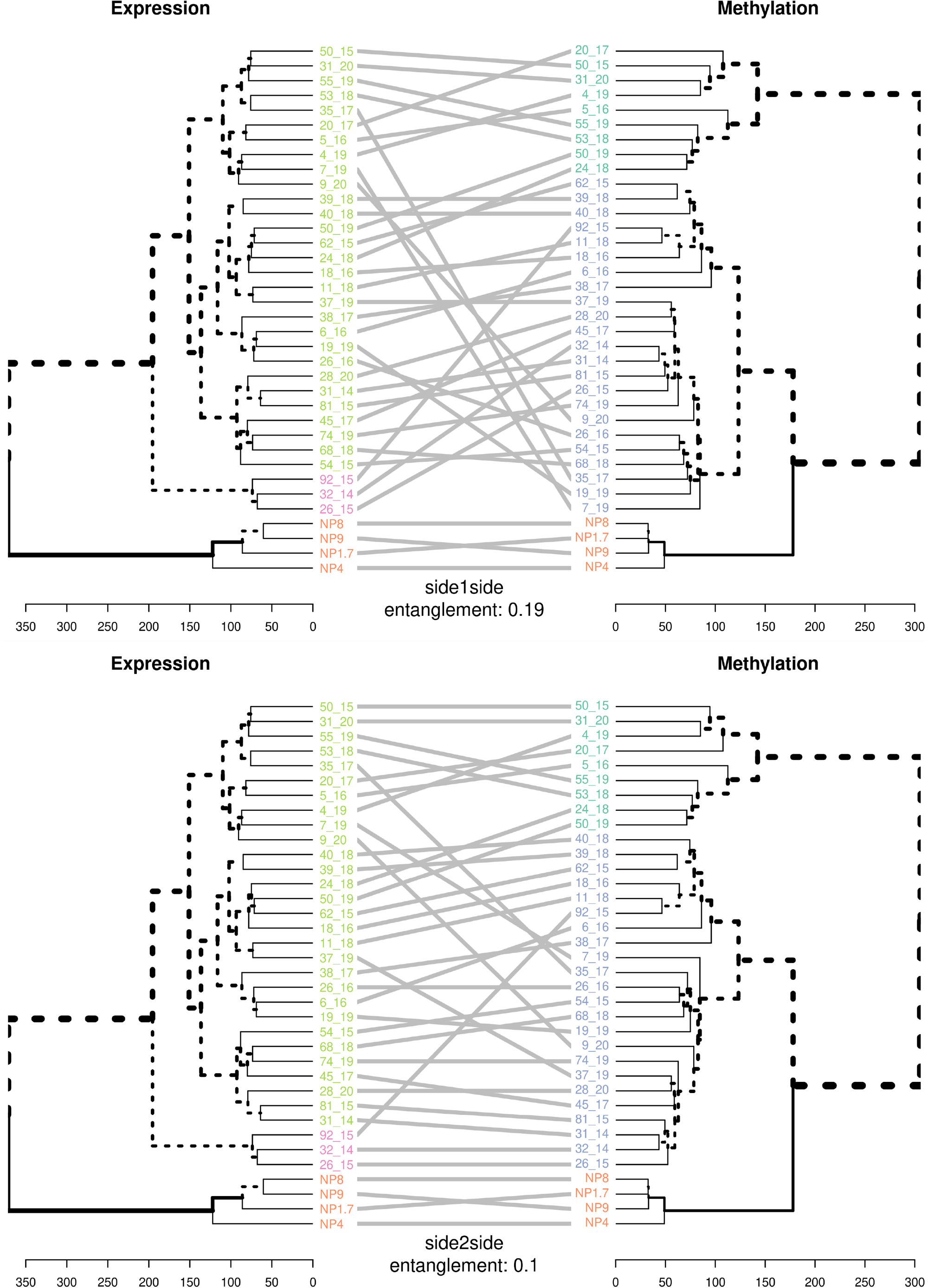

Furthermore, chordoma microenvironment can be analyzed in the wider context of other neoplasms. Data from The Cancer Genome Atlas, deconvoluted with ESTIMATE method were downloaded and scores generated by ESTIMATE method on our expression data were compared to them. Both stromal and immune scores were significantly higher in chordoma I than in chordoma C (Mann-Whitney p: 6e-5 and 0.022, respectively). Comparison with other cancer types revealed amounts of immune infiltrate in chordomas I comparable to glioblastoma, bladder and colorectal cancer, while chordomas C have median immune score remarkably lower than other human cancers, as visualized in Figure 4d.

### DNA copy number changes in two epigenetic subtypes of chordomas

EPIC DNA methylation array allows for the evaluation of large, unbalanced, structural chromosomal genomic amplifications and deletions (commonly referred to as Copy Number Variations, CNVs). Copy number analysis was applied to all available chordoma samples and with a cut-off of 0.3 for absolute copy number change; 261 copy number events were identified in all samples (Figure 5a). The mean number of segments was 7.19 with quantile distribution of [0, 1, 2.5, 11.25, 34]. Moreover, CNVs were more common in chordoma C cluster as compared to chordoma I (median 14 vs 1, p = 0.02).

The main difference in CNVs between two chordoma clusters were chromosomic losses in chromosome 9 region, containing *CDKN2A/B*, affecting chordoma C cluster. This loss was found in 9 of 10 tumors from chordoma C cluster while it was observed only in 1 out of 23 tumors of cluster I, with CNV score cut-off at 0.3 (exact Fisher test p = 7.95e-5). This difference is also significant, when comparing conumee scores for this region (Figure 5b). In order to verify the functional consequences of chromosome 9 deletions, expression levels of genes localized in commonly deleted regions were examined. Expression of 9p21-located genes, containing *CDKN2A*, differentiated the clusters most significantly (Figure 5c and 5d, Supplementary Figure 3). As mentioned earlier, *CDKN2A* was overexpressed in chordoma I cluster (Figure 5e).

### Prognostic factors in chordomas

Various features, including methylation cluster, immune infiltration, WGCNA modules, and *CDKN2A* loss were tested for effect on overall survival. However, only the total number of CNVs showed a significant association with survival – patients with more variations had a higher risk of death (Figure 5e). This effect was seen in both uni- and multivariate analysis (Table 3).

## Discussion

Our study aimed to determine DNA methylation profile in skull base chordomas as well as to investigate their possible methylation-based subclassification and the role of epigenetic abnormalities in their pathogenesis. The results indicate that there are two molecular subtypes of these tumors. These subtypes have distinct DNA methylation profile reflected by difference in global methylation level and several differentially methylated regions. Accordingly, two epigenetic – DNA methylation-based subtypes of chordomas were also identified in two very recently published studies [13,14]. In contrast with these studies, we have also characterized these subtypes by comparison with normal DNA methylation patterns obtained from NP. When analyzing global methylation pattern, we found chordoma I to be quite similar to control samples with comparable overall DNA methylation level and hypermethylation of 5’ flanking regions of the genes and CGIs, while the other subtype turned out to be generally hypomethylated with an increased methylation level at CGIs. This epigenetic landscape of whole genome hypomethylation with hypermethylated CGIs resemble general DNA-methylation profile of cancer cells [41,42]. A large number of aberrantly methylated regions was identified in each chordoma subtype. Comparison of chordomas vs NP showed DMRs in many cancer-related genes (Figure 1e) as well as in genomic cluster encoding homeobox domain genes (at *HOXD3, HOXD4* and *HOX11, HOXA9, HOXA10*) that play an important role in both normal differentiation and tumorigenesis [43]. Comparison of two chordoma subtypes showed DMRs in loci encoding genes with a known role in tumorigenesis but also in regions covering numerous genes encoding short RNAs (micro RNAs and small nucleolar RNA (snoRNAs)). Both classes of small RNA play a role in cancer but their significance in pathogenesis of chordomas is poorly recognized (especially for snoRNAs) [44,45] and our results are the first that indicate epigenetic misregulation of small RNAs in these tumors.

RNA sequencing of the same tumor samples that were included in methylation analysis allowed for further characterization of the methylation subtypes, beyond previous analysis [13,14]. Clustering based on gene expression did not show a clear overlap between expression and methylation-based classification of the samples and most of chordoma samples were classified in the same expression cluster. No gene expression-based subclassification of skull base chordomas was also found in previous RNAseq analyses [16,46]. The existence of 3 distinct expression groups of skull base chordomas was described recently [18], however, it was not observed in our data.

Correlation-based analysis of the role of DNA methylation in gene expression suggests that in chordomas only a small proportion of all the genes is-controlled by promoter DNA methylation. However, the fact that genes with promoter methylation/expression correlation are enriched for the genes that are differentially expressed in chordomas indicates that DNA methylation contributes to pathogenesis of these tumors.

The analysis of the role of DNA methylation level at the identified DMRs in the expression of the genes located at these regions reveals that only minor proportion of DMRs has a role in differential gene expression. These expression change-related DMRs include some genes that are known to play a role in tumorigenesis, including *PIK3CD*,[47] *UNC5D*, [48] or *NMU* [49].

Interestingly, looking for more general relationship between promoter methylation and the expression of genes in chordoma samples we found brachyury gene expression to be correlated with DNA methylation. Lower promoter methylation was related to higher *TBXT* expression in chordomas that is in line with the results on the regulation of this gene during cell differentiation. Lowering brachyury expression was associated with differentiation of mesenchymal stem cells into osteoblasts and adipocytes and it was accompanied by progressive methylation of its promoter region [50]. Previous observations in chordomas suggested rather a role of histone than DNA methylation in epigenetic regulation of *TBXT* [51]. We assume that both elements of epigenetic regulation may contribute to upregulating brachyury expression in chordomas.

Transcriptomic profiling in chordomas allowed for functional analysis of the differences in gene expression between cluster C and cluster I tumors. The results clearly demonstrated that distinct molecular processes are involved the pathogenic mechanism of the two tumor subtypes. Cluster I tumors have gene expression pattern that indicates high immune infiltration, while cluster C tumors appear to be driven by the processes related to mitosis. Additionally, pathway analysis of genes differentially expressed between two chordoma subtypes that are controlled by DNA methylation (as examined with correlation analysis) showed overrepresentation of genes related mainly to immune infiltration pathways. Enrichment in immune component in chordoma I was therefore observed independently in both gene expression analysis and the analysis of the relationship between DNA methylation and expression levels. These observations were confirmed by a downstream analysis with deconvolution methods, and copy number analysis. Estimation based on methylation arrays, RNA-seq and anti-CD8 immunostaining of tumor tissue samples consistently show that chordoma cluster I tumors are highly enriched in immune cells. Comparable results were observed in studies by Huo X et al. and Zuccato JA et al. Both groups reported that one of DNA methylation chordoma subtypes has notably higher content of immune cells [13,14]. Using gene expression data, we compared the estimated immune infiltration status in chordomas and common human cancers. This clearly highlighted the difference between subtypes and showed that the so-called “immune cold” chordoma C subtype generally has immune scores below those observed in other caners (Figure 4d).

We used data generated with EPIC microarrays for DNA copy number analysis. It showed that, in contrary to chordoma I, chordoma C subtype is notably affected by DNA copy number changes with deletion of 9p chromosomal arm as most common aberration found in nearly all tumors in this methylation cluster. This chromosomal arm contains *CDKN2A* and *CDKN2B* genes - crucial cyclin-dependent kinase inhibitors. Loss of these key cell cycle suppressors in C-type chordomas clearly corresponds to results of gene set enrichment analysis which revealed the role of cell proliferation in the pathogenesis of this chordoma subtype. Our result showing the occurrence of 9p deletion in nearly all tumors of C subtype slightly differs from the observation by Huo X et al and Zuccato JA et al. Both groups also showed that CNVs are more frequent in one of methylation subtypes of chordoma but they identified 9p loss only in a minor proportion of specimens. However, Huo X et al reported a frequent loss of chromosome arm 19p (encoding cyclin E gene) in immune enriched subtype of chordomas that was found neither in our results nor in those published by Zuccato JA et al. Some differences in molecular features of the tumors as well as slightly different abundance proportions between two subtypes that are observed in ours and previously reported studies may by caused by populational differences. Previous studies included Canadian/French [13] and Chinese [14] patients while our cohort is composed of Polish patients only.

A notable effort in chordoma research is focused on identification of prognostic factors. DNA methylation-based subclassification of skull base chordomas showed a clinical relevance. Both the results by Zuccato JA et al and Huo X et al indicated prognostic role of immune component in DNA methylation pattern, but the authors found dissimilar results. Observations by Zuccato JA indicate shorter overall survival in patients with immune-hot chordoma subtype identified in DNA methylation-based clustering. In contrary, Huo X et al reported worse outcome in patients with low immune cell content and higher tumor purity. When we compared overall survival between two DNA methylation subtypes of chordomas we did not find significant difference between two methylation clusters. The molecular features that we found as discriminating epigenetic chordoma subtypes are known as relevant prognostic factors in other studies. Higher immune infiltration status determined by microscopic assessment of CD3+ and CD8+ cells count was identified as related to better prognosis in patients with spinal chordomas [52,53] but another study showed that higher content of CD8-positive cells is related to shorter survival [54].

In previous studies, DNA copy number status in skull base chordomas also showed prognostic value. Deletion of 9p21, that we observed in nearly all subtype C chordomas was associated with worse recurrence-free survival but not OS in study by ai J et al[10]. The same study also showed a prognostic relevance of 22q (encoding *SMARCB1* gene) deletions [10]. Changes in 22q region were not observed in our patients’ group probably because it is more homogeneous and contained only one histological subtype of classical chordomas. Clinical relevance of CDKN2A expression and deletion of this gene locus were specifically address in the other study on classical and chondroid chordomas [39]. It revealed *CDKN2A* deletion in approximately 50% of the tumors but significant relationship between DNA copy number and patients survival was not observed [39].

Our results of copy number analysis showed prognostic value of chromosomal instability. We did not find survival difference between patients with 9p (*CDKN2A* locus) loss and 9p stable patients, but we observed a significant relationship of the higher level of copy number changes in tumors with shorter survival as clearly illustrated by significant difference in OS between patients with high and low number of CNVs. This observation is concordant with a general finding that chromosomal instability in human cancer is a biomarker of poor prognosis [55]

The results showing prognostic significance of clustering chordoma samples using transcriptomic data were also recently published. Bai J et al identified three gene expression-related clusters of chordomas and subsequent comparison of two of these clusters showed a difference in patients’ OS. Their observations were not confirmed by our analysis since we did not observe such clustering of tumors with RNA-seq results. Most of the tumors were grouped in the same major cluster and we did not find relationship between transcriptomic subtype and OS.

## SUMMARY

Two distinct chordoma subtypes (subtype C and I) with different patterns of aberrant DNA methylation have been identified upon genome-wide DNA methylation analysis. They have different profile of both global and locus-specific methylation pattern as well as distinct gene expression. Differences in gene expression indicate immune activation in I chordomas and enhanced cell proliferation in C chordomas. Immune enrichment in I chordomas found in transcriptomic profiling is also confirmed by results of analysis with deconvolution methods. C chordomas are characterized by higher chromosomal instability, according to results of copy number analysis and they have 9p deletion causing downregulation of cell cycle inhibitors *CDKN2A/B*. Tumor subtypes do not differ significantly in terms of prognosis, however, there is a significant influence of chromosomal instability level on shorter survival.

## Supporting information

Tables and figures descriptions

## List of abbreviations

5’UTR: 5’ untranslated region
Bp: base pairs
CGI: CpG island
CNVs: Copy Number Variations
DEG: Differentially expressed gene
DMP: differentially methylated probe
DMR: differentially methylated region
GO: Gene Ontology
GSEA: Gene set enrichment analysis
NP: nucleus pulposus
RNA-seq: RNA sequencing
snoRNAs: small nucleolar RNA
SNP: single nucleotide polymorphism
TSS: transcription start site
WHO: World Health Organization
WGCNA: weighted correlation network analysis

## Declarations

### Ethics approval and consent to participate

The study was conducted in accordance with the Declaration of Helsinki, and approved by the Institutional Ethics Committee of Maria Sklodowska-Curie Institute - Oncology Center in Warsaw, Poland (approval decision 32/2020, dated 18.06.2020). Informed consent for the use of tissue samples was obtained from all subjects involved in the study.

### Consent for publication

Not applicable

### Availability of data and material

The datasets generated with RNAseq and EPIC arrays aare available at Gene Expression Omnibus repository, GSE230168 (https://www.ncbi.nlm.nih.gov/geo/query/acc.cgi?acc=GSE230168).

### Authors’ contributions

Szymon Baluszek: Conceptualization, Methodology, Formal analysis, Investigation, Visualization, Writing— review and editing; Paulina Kober: Formal analysis, Investigation, Writing—original draft preparation; Natalia Rusetska: Investigation; Jacek Kunicki: Resources, Data curation; Tomasz Mandat: Resources, Data curation, Conceptualization, Funding acquisition; Mateusz Bujko: Conceptualization, Methodology, Formal analysis, Visualization, Writing—original draft preparation

## Acknowledgements

Not applicable

