## Supplementary material for "DNA methylation, combined with RNA sequencing, provide novel insight into molecular classification of chordomas and their microenvironment": Tables and figures descriptions

### Figures and tables

#### Table 1

Characteristics of skull base chordoma patients.

| | |
|---|---|
| Number of patients | n=32 |
| Sex<br>Females<br>Males | <br>15/32 (47%)<br>17/32 (53%) |
| Age (years; median (range)) | 60 (23-76) |
| Skul base location<br>Clivus<br>Clisvus and craniovertebral junction | <br>21/32 (65,6%)<br>11/32 (34,4%0 |
| Surgery type<br>Endoscopic endonasal<br>craniotomy | <br>31/32 (96,9%)<br>1/32 (3,3%) |
| Gross resection rate<br>Complete<br>Subtotal<br>Partial | <br>11/32 (34,4%)<br>16/32 (50%)<br>5/32 (15,6%) |
| Recurrence status<br>Newly diagnosed<br>Recurrent | <br>22 /32 (69%)<br>10/32 (31%) |
| Tumor size - max. diameter (mm; median (range)) | 8,5 (1-17) |
| Histological type<br>Classical chordoma | <br>32 (100%) |
| Death status<br>No<br>Yes<br>Follow up (months - median (range)) | <br>19/32 (59%)<br>13/32 (41%)<br>38 (6-97) |

#### Table 2

The numbers of differentially methylate probes and regions as wells as differentially expressed genes for each chordoma of nucleus pulposus sample group

| First group | I chord. | C chord. | I chord. | NP | C chord. | NP | chord. | NP |
|---|---|---|---|---|---|---|---|---|
| Second group | C chord. | I chord. | NP | I chord. | NP | C chord. | NP | chord. |
| DMPs | 4892 | 9 | 22676 | 1261 | 1532 | 1502 | 18064 | 1383 |
| DMRs | 13622 | 180 | 15705 | 1208 | 2825 | 3245 | 11670 | 1083 |
| DEGs | 1128 | 170 | 5078 | 4052 | 3426 | 3426 | 5293 | 4252 |
| WGCNA modules | 4 | 0 | 12 | 11 | 8 | 9 | 13 | 13 |

#### Table 3

Survival effect of selected clinical and molecular features in a uni- and multivariate Cox hazard model

| Variable | Trait | Multivariate | | | Univariate | | |
|---|---|---|---|---|---|---|---|
| | | Coefficient | Z | p-value | Coefficient | Z | p-value |
| CNVs | | 0.16 | 2.92 | 0.003 | 0.08657 | 3.05 | 0.002 |
| Immune infiltrate | | 6.70 | 1.60 | 0.11 | -2.32 | -1.06 | 0.29 |
| Sex | male | -1.12 | -1.45 | 0.15 | 0.05 | 1.84 | 0.07 |
| Age | | 0.04 | 1.40 | 0.16 | -0.45 | -0.77 | 0.44 |
| Cluster | I chord. | 0.60 | 0.61 | 0.54 | -0.32 | -0.52 | 0.60 |
| | | Likelihood ratio test=15.56 on 5 degrees of freedom, p=0.008 events: 13 of 31 observations | | | | | |

#### Figure 1

Results from EPIC DNA methylation arrays a) Heatmap of scaled methylation M-values of 3648 most variable probes, split in rows, according to CpG relation to CGI and clustering of samples. b) Beta values of 364,784 probes with standard deviation of β-values above 0.1, split according to CpG relation to CGI and gene. c) Quantification of differences in overall methylation levels using a linear model of M-values. d) Distribution of differentially methylated probes classified according to their position regarding CGI and genes. e) Differentially methylated regions, depicted on Manhattan plots (genomic position on x-axis), gene names for most significant and biologically interesting DMRs are captioned.

#### Figure 2

Analysis of genes expression and its relation to the DNA methylation profile. a) Heatmap of 4275 top most variably expressed genes and their clustering showing lack of clear overlap between methylation and expression-based chordoma clusters. b) Fraction of genes differentially expressed between chordoma subtypes as well as chordomas and nucleus pulposus that are under DNA methylation control (as defined by significant correlation of mean promoter methylation with gene expression), the blue dot-and-dash line indicates the level of methylation control of the genome in general c) Volcano plot of differentially expressed genes identified in chordoma – nucleus pulposus comparison d) Volcano plot of differentially expressed genes found in chordoma I – chordoma C comparison e) Alluvial plot of DMR relation to DEGs in chordoma I – chordoma C comparison, height of the bars represents number of genes/DMRs f) Role of DNA methylation in *TBXT* (brachyury) gene. Difference in the methylation levels of CpGs at *TBXT* locus with DMPs labeled with * (for adj.p<…) or *** (for adj.p<…) (left panel), difference in *TBXT* expression of (middle panel) and correlation between *TBXT* promoter methylation and expression levels (right panel); p-values are shown for chordoma-nucleus pulposus comparison g) Difference in the methylation levels of CpGs at *PTPRCAP* locus with DMPs labeled with * (for adj.p<…) or *** (for adj.p<…) (left panel), difference in *PTPRCAP* expression of (middle panel) and correlation between *PTPRCAP* promoter methylation and expression levels (right panel) h) Position of *TBXT* (brachyury) and *PTPRCAP* genes on their chromosomes, along with EPIC methylation probes position and DMRs.

#### Figure 3

Functional analysis of the RNA sequencing results a) Top 20 terms of gene set enrichment analysis on gene ontology terms (chordoma I – chordoma C comparison) b) Differentially expressed genes and terms from gene set enrichment analysis. Due to number of terms and genes top hits, basing on p-value, were selected

c) Weighted gene co-expression network analysis, presented on heatmap; samples are shown in columns, WGCNA modules in rows, significance of tests is shown in the panel on the left d) WGCNA module 3 (with significantly higher score for chordoma I samples) intersected with STRING database; genes to plot were selected based on top connectivity (importance for the module) and centrality (best connected within module); gene names in bold were also differentially expressed in chordoma I, compared with chordoma C e) Analogous for module 2, higher in chordoma versus nucleus pulposus, DEGs for chordoma – NP comparison are shown in bold.

#### Figure 4

Cell type deconvolution of RNA-seq and DNA methylation data a) heatmap presenting scores from two independent methods – MethylResolver (utilizing EPIC DNA methylation data, upper panel) and MCPcounter (utilizing RNA-seq data, lower panel) both point to higher immune infiltration in chordoma I cluster, significant differences in U-Mann-Whitney test are marked with * b) Correlation matrix of immune signatures from both methods (Kendall correlation); coefficients are shown in the middle of each cell, significant ones are black c) Plot presenting signature correlation for cytotoxic T-lymphocytes from both methods, correlation coefficient: 0.73, adjusted p-value: 2.8e-142, cluster differences adjusted p-values: 0.0012 and 0.0003 for MethylResolver and MCPcounter, respectively d) Comparison of chordomas (including chordoma subtypes classified according to DNA methylation profile) with existing signatures for variable human cancer types from ESTIMATE method (signatures for immune and stromal components of the tumor).

#### Figure 5

Estimations of copy number alterations and their biological and prognostic relevance a) Results of copy number variation (CNV) imputation from EPIC DNA methylation array performed by conumee method. Genomic segments with absolute scores above 0.3 are shown. Sample label colors correspond to methylation clusters b) Box plot for conumee score of *CDKN2A* gene locus; 0.3 cut-off marked with dashed line c) Estimation of gene expression across selected regions of chromosome 9 (according to chromosomal bands), calculated by GSEAlm package; label colors correspond to methylation clusters d) Box plot of GSEAlm score for chr9p21 band (containing *CDKN2A*/*B* genes*)* that has lowest p-value as the result of comparing two chordoma subtypes in terms of the expression across the whole genome. e) *CDKN2A* expression across methylation clusters and normal nucleus pulposus f) Kaplan–Meier plot for number of CNVs (0.3 cut-off in conumee estimate); Cox proportional hazard model was used for testing, cut-off of 8 CNVs was picked by maxstat package for visualization purposes.

#### Supplementary tables and file

Can be found here (due to BioArchives limit): tiny.cc/chor_sf_bioarch_2023
